# The tumor suppressor BRCA1/BARD1 complex localizes to the synaptonemal complex and regulates recombination under meiotic dysfunction in *Caenorhabditis elegans*

**DOI:** 10.1101/280909

**Authors:** Qianyan Li, Takamune T. Saito, Alison J. Deshong, Marina Martinez Garcia, Saravanapriah Nadarajan, Katherine S. Lawrence, Paula M. Checchi, Monica P. Colaiacovo, JoAnne Engebrecht

## Abstract

Breast cancer susceptibility gene 1(BRCA1) and binding partner BRCA1-associated RING domain protein 1 (BARD1) form an essential E3 ubiquitin ligase important for DNA damage repair and homologous recombination. In *Caenorhabditis elegans* BRCA1/BRC-1 and BARD1/BRD-1 orthologs are not essential, but function in DNA damage repair and homologous recombination, as well as in meiosis. In proliferating germ cells and in early meiotic prophase, BRC-1 and BRD-1 are nucleoplasmic, with enrichment at foci that partially overlap with the recombinase RAD-51. In mid-pachytene, BRC-1 and BRD-1 are observed on tracks, before concentrating to the short arms of bivalents, co-localizing with a central region component of the synaptonemal complex. We found that BRD-1 is essential for BRC-1 to associate with chromatin and the synaptonemal complex, but BRC-1 is not required for BRD-1 localization; the complex fails to properly localize in the absence of either meiotic recombination or chromosome synapsis. Inactivation of BRC-1/BRD-1 enhances the embryonic lethality of mutants that perturb chromosome synapsis and crossover recombination, suggesting that BRC-1/BRD-1 plays an important role in monitoring recombination in the context of the synaptonemal complex. We discovered that BRC-1/BRD-1 stabilizes the RAD51 filament when the formation of a crossover-intermediate is disrupted. Further, in the absence of BRC-1/BRD-1 crossover distribution is altered, and under meiotic dysfunction, crossover numbers are perturbed. Together, our studies indicate that BRC-1/BRD-1 localizes to the synaptonemal complex where it serves a checkpoint function to monitor and modulate meiotic recombination.

**Project Summary:** Our genomes are passed down from one generation to the next through the specialized cell division program of meiosis. Meiosis is highly regulated to coordinate both the large scale chromosomal and fine scale DNA events to ensure fidelity. We analyzed the role of the tumor suppressor BRCA1/BARD1 complex in meiosis in the worm, *Caenorhabditis elegans*. We find that BRCA1/BARD1 localizes dynamically to the proteinaeous structure that aligns maternal and paternal chromosomes, where it regulates crossover recombination. Although BRCA1/BARD1 mutants have only subtle meiotic defects, we show that this complex plays a critical role in meiotic recombination when meiosis is perturbed. These results highlight the complexity of ensuring accurate transmission of the genome and uncover the requirement for this conserved complex in meiosis. As women carrying BRCA1 mutations with no indication of cancer have fertility defects, our results provide insight into why BRCA1 mutations impact reproductive success.

## Introduction

BRCA1 was identified twenty-eight years ago as the causative agent of early-onset familial breast cancer (1). Subsequently, BRCA1 was shown to interact with BARD1 through their RING domains (2), to form an E3 ubiquitin ligase, which adds the small polypeptide ubiquitin to protein substrates (3). While BRCA1/BARD1 has been extensively studied with respect to its crucial tumor suppressor activities, we still do not fully understand how this protein complex mediates the diverse functions that have been ascribed to it [*e.g.*, DNA metabolism, checkpoint signaling, chromatin dynamics, centrosome amplification, and transcriptional and translational regulation (4, 5)]. This is due in part to the diversity of protein-protein interactions involved in generating numerous distinct BRCA1/BARD1 complexes (6). An additional impediment to understanding BRCA1/BARD1 function is that the corresponding mouse knockouts are embryonic lethal (7, 8).

The simple metazoan *Caenorhabditis elegans* offers several advantages to the study of this key complex. First, unlike in mammals, *C. elegans* BRCA1 and BARD1 orthologs, BRC-1 and BRD-1, are not essential yet play critical roles in DNA replication and the DNA damage response, as well as in homologous recombination, which is essential for repairing programmed double strand breaks (DSBs) during meiosis (9-14). Additionally, attributes of the *C. elegans* system, including sophisticated genetics, ease of genome editing, and the spatio-temporal organization of the germ line allows us to overcome some challenges inherent in studying this complex in mammalian meiosis.

Meiosis is essential for sexual reproduction and results in the halving of the genome for packaging into gametes. During meiosis, homologous chromosomes are connected by crossover recombination to facilitate their alignment and segregation on the meiotic spindle. Recombination is integrated and reinforced with chromosome pairing and synapsis, although the extent of dependencies of these critical meiotic processes are distinct in different organisms (reviewed in (15, 16). While it is well established that BRCA1 plays an important role in DNA repair and recombination (5), the specific function of BRCA1/BARD1 in meiotic recombination is not known. In mice, partial deletions of BRCA1 result in early apoptosis due to failures in meiotic sex chromosome inactivation (17, 18). BRCA1 has been shown to co-localize with RAD51 on asynapsed chromosomes in mouse spermatocytes, suggesting it functions in meiotic recombination (19). In *C. elegans, brc-1* mutants have mild meiotic phenotypes consistent with a role in some aspect of meiotic recombination (9, 10). However, the relationship between BRC-1/BRD-1 function in synapsis and recombination has not been explored.

Here, we assessed BRC-1 and BRD-1 dynamics in the *C. elegans* germ line. Surprisingly, BRC-1/BRD-1 localizes to the synaptonemal complex (SC), becomes concentrated onto chromosome regions upon crossover designation, and at late meiotic prophase is restricted to the short arm of the bivalent. We found that BRC-1/BRD-1 concentration at late pachytene precedes SC reorganization around the crossover site. Further, our data reveal a role for the BRC-1/BRD-1 complex in promoting homologous recombination by protecting the RAD-51 filament and altering recombination outcomes under meiotic dysfunction. Similar findings are reported by Janisiw et al. in the accompanying paper.

## RESULTS

### GFP::BRC-1 and BRD-1::GFP are expressed in embryos and the germ line

To examine BRC-1 and BRD-1 expression and localization in *C. elegans*, we engineered GFP::BRC-1 and BRD-1::GFP fusions at the endogenous loci using CRISPR/Cas9 (20). *brc-1* and *brd-1* mutants produce slightly elevated levels of male progeny (X0), a readout of *X* chromosome nondisjunction, have low levels of embryonic lethality and display sensitivity to γ-irradiation (IR) (10). Worms expressing these fusions as the only source of BRC-1 or BRD-1 produce wild-type levels of male progeny and embryonic lethality and are not sensitive to IR (S1 Fig), indicating that the fusions are fully functional.

We monitored the localization of GFP::BRC-1 and BRD-1::GFP by live cell imaging. In whole worms, GFP fluorescence was observed in embryos and throughout the germ line, with very little signal in the soma (note auto-fluorescence of gut granules also observed in wild type; Fig 1A). Immunoblots of whole worm extracts of *gfp::brc-1; fog-2*, which are true females (21) and therefore do not contain embryos, compared to self-fertilizing *gfp::brc-1* hermaphrodites containing embryos, revealed that <10% of the GFP::BRC-1 signal is due to expression in embryos (S1 Fig). Thus, BRC-1/BRD-1 is expressed at significant levels throughout the germ line, consistent with a role of this complex in both mitotically-dividing germ cells and in meiosis (9-12, 14).

**Fig 1.**
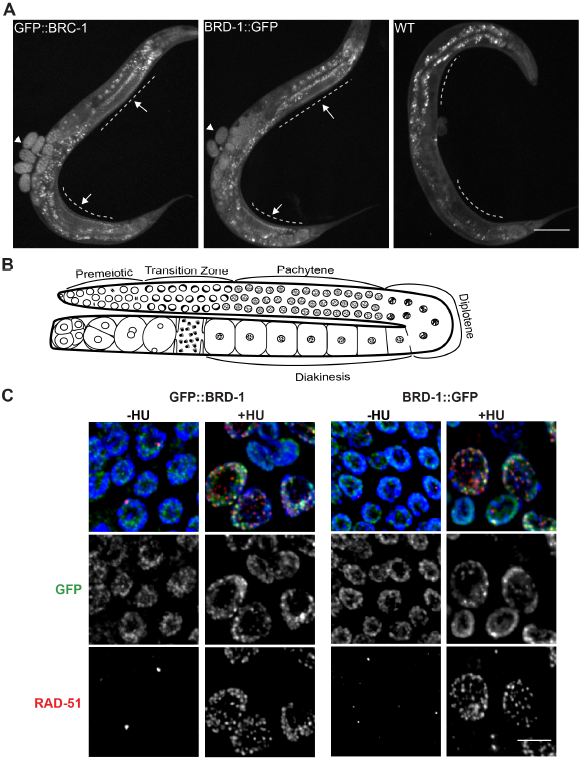
GFP::BRC-1 and BRD-1::GFP are predominately expressed in the germ line and respond to stalled replication forks. A) GFP fluorescence of whole worms expressing GFP::BRC-1, BRD-1::GFP, or no GFP (WT). Dashed line denotes germ line with arrows to indicate GFP fluorescence; arrowheads denote GFP signal in embryos; gut granules auto-fluoresce. Scale bar = 100μm. B) Schematic of the spatiotemporal organization of the hermaphrodite germline with meiotic stages indicated. C) Proliferating germ cells expressing GFP::BRC-1 or BRD-1::GFP (green), stained with antibodies against RAD-51 (red), and counterstained with DAPI (blue) in the absence (-HU) and presence of 5mM hydroxyurea (+HU) (D). Scale bar = 10μm.

### GFP::BRC-1 and BRD-1::GFP are enriched at stalled/collapsed replication forks in proliferating germ cells

The *C. elegans* germ line is arranged in a spatio-temporal gradient, with proliferating germ cells and all stages of meiosis arrayed from the distal to proximal end (22) (Fig 1B). In proliferating germ cells, GFP::BRC-1 and BRD-1::GFP were nucleoplasmic, with regions of brighter fluorescence. There was partial co-localization with the RAD-51 recombinase, which marks stalled replication forks (23) (Fig 1C). Work in both mammalian cells and *C. elegans* have revealed that BRCA1 mediates repair of stalled/collapsed replication forks (14, 24), and in mammals this function is independent of the established role of BRCA1 in homologous recombination (25). To determine whether BRC-1/BRD-1 concentrates at stalled or collapsed replication forks, we treated worms with the ribonucleotide reductase inhibitor, hydroxyurea (HU). HU slows replication causing fork stalling and collapse, and cell cycle arrest leading to enlarged nuclei (23, 26). GFP::BRC-1 and BRD-1::GFP fluorescence became more punctate following exposure to HU and ~50% of these regions co-localized with RAD-51 (Fig 1C). Consistent with a role in resolving collapsed replication forks, both *brc-1* and *brd-1* mutants were sensitive to 5mM HU as measured by embryonic lethality (S1 Fig). We also observed more GFP::BRC-1 foci, which partially overlapped with RAD-51, following IR treatment (S1 Fig), as has been previously reported for BRD-1 (27). Together, these results suggest that BRC-1/BRD-1 responds to and concentrates at stalled/collapsed replication forks, and IR-induced lesions. As there is not a one-to-one correspondence with RAD-51, BRC-1/BRD-1 appears to only transiently associate with RAD-51.

### GFP::BRC-1 and BRD-1::GFP localize to the SC and concentrate to the short arm of the bivalent during meiotic prophase

In early meiotic prophase (transition zone/early pachytene), GFP::BRC-1 and BRD-1::GFP were observed diffusely on chromatin (Fig 2A). Regions of more intense GFP::BRC-1/BRD-1::GFP fluorescence partially overlapped with RAD-51, which marks meiotic DSBs (28). Beginning at mid-pachytene, GFP::BRC-1 and BRD-1::GFP were observed in tracks along the entire chromosome length, and then concentrated to a portion of each chromosome at late pachytene (Fig 2A). In diplotene/diakinesis, GFP::BRC-1 and BRD-1::GFP were further restricted to six short stretches, corresponding to the six pairs of homologous chromosomes. As oocytes continued to mature, GFP::BRC-1 and BRD-1::GFP were disassembled from chromosomes in an asynchronous manner, with some chromosomes losing signal before others. Thus, in diakinesis nuclei we did not always observe six stretches of fluorescence, and the fluorescence intensity varied between chromosomes.

**Fig 2.**
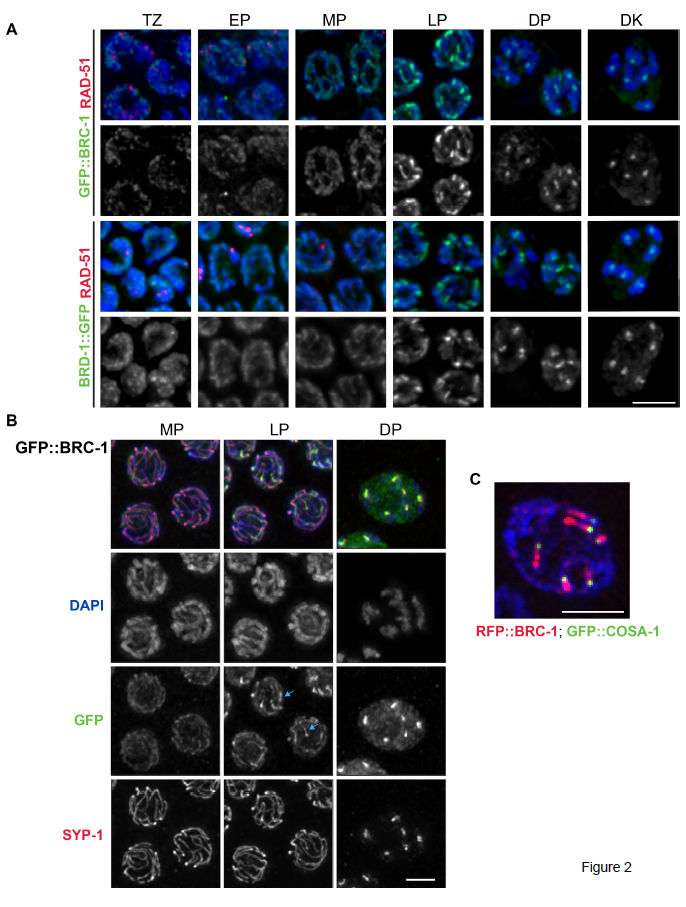
GFP::BRC-1 and BRD-1::GFP localize to the SC in meiotic prophase. A) Nuclei from indicated meiotic stages stained with RAD-51 antibodies (red), DAPI (blue) and imaged for GFP fluorescence (green). TZ = transition zone; EP = early pachytene; MP = mid pachytene; LP = late pachytene; DP = diplotene; DK = diakinesis. Scale bar = 5 μm. B) Co-localization between GFP::BRC-1 (green) and SC central component SYP-1 (red); germ lines at indicated stages were counterstained with DAPI. Blue arrows show chromosomal regions where GFP::BRC-1 concentrates before SYP-1. Scale bar = 2 μm. C) RFP::BRC-1 (red) and GFP::COSA-1 (green) at late pachytene showing RFP::BRC-1 on one side of the GFP::COSA-1 foci, which marks the persumptive crossover. Scale bar = 2 μm.

Co-staining with antibodies against the SC central region component SYP-1 (29), revealed that the BRC-1/BRD-1 tracks correspond to the SC (Fig 2B). Interestingly, the concentration of GFP::BRC-1 to a portion of each chromosome precedes the relocalization of SYP-1 (arrows in late pachytene images of GFP::BRC-1 and SYP-1; Fig 2B). As the SC reorganizes as a consequence of crossover maturation (30), we examined worms co-expressing RFP::BRC-1 and GFP::COSA-1, a cyclin related protein that marks presumptive crossover sites (31). RFP::BRC-1 is also fully functional (S1 Fig), although the fluorescent signal is weaker than GFP, and could only be detected in mid-late pachytene through diakinesis. GFP::COSA-1 was observed at one end of each RFP::BRC-1 stretch (Fig 2C). Thus, BRC-1 and BRD-1 partially associate with RAD-51 in proliferating germ cells and early in meiosis but beginning in mid pachytene, GFP::BRC-1 and BRD-1::GFP co-localize with the SC. BRC-1/BRD-1 eventually retracts to the short arm of the bivalent and its concentration to a portion of each chromosome precedes SC reorganization around the crossover site, as marked by COSA-1. These results are consistent with BRC-1/BRD-1 functioning in one or more aspects of meiotic recombination within the context of the SC.

### BRD-1 is required for GFP::BRC-1 localization but BRC-1 is not required for BRD-1::GFP localization

In both mammalian cells and *C. elegans*, BRCA1/BRC-1 and BARD1/BRD-1 form a stable complex (2, 27). To probe the relationship between *C. elegans* BRC-1 and BRD-1 *in vivo*, we imaged live worms heterozygous for both RFP::BRC-1 and BRD-1::GFP (*brc-1* and *brd-1* are linked). In the heterozygous state the RFP signal could only be detected at late pachytene through early diakinesis when BRC-1 and BRD-1 are concentrated on short tracks. The RFP and GFP signals overlapped, indicating that BRC-1 and BRD-1 are localized together on the SC (Fig 3A).

**Fig 3.**
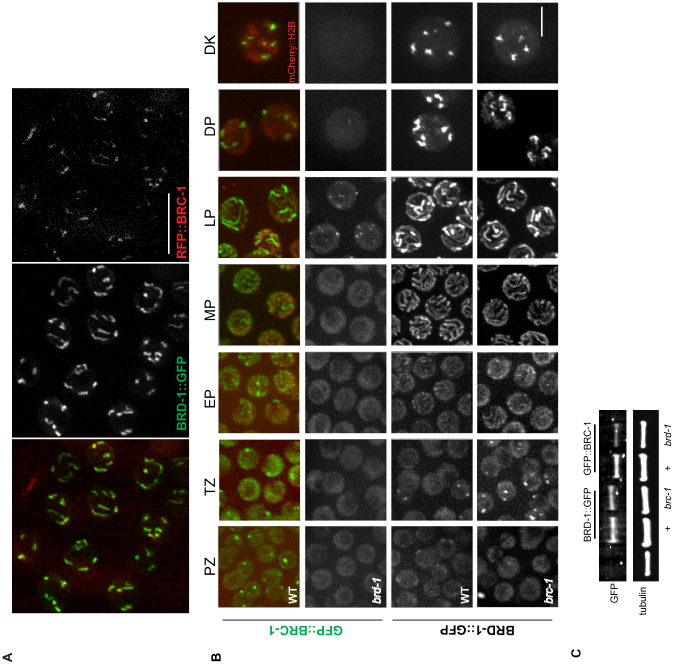
BRD-1 is required for BRC-1 localization but BRC-1 is not required for BRD-1 localization. A) Co-localization between BRD-1::GFP (green) and RFP::BRC-1 (red) at late pachytene in live worms. Scale bar = 10 μm. B) Stills of germline nuclei from live worms expressing GFP::BRC-1 and mCherry:Histone H2B (WT; top panel) and GFP::BRC-1 expression in the *brd-1* mutant at indicated meiotic stages. Bottom two panels show BRD-1::GFP localization in wild type and the *brc-1* mutant. TZ = transition zone; EP = early pachytene; MP = mid pachytene; LP = late pachytene; DP = diplotene; DK = diakinesis. Scale bar = 5 μm. C) Immunoblot of whole worm extracts from indicated worms probed with anti-GFP and *α*-tubulin antibodies, as loading control.

To examine localization dependencies between BRC-1 and BRD-1 in *C. elegans* germ cells, we monitored GFP::BRC-1 and BRD-1::GFP in the corresponding *brc-1* and *brd-1* mutant backgrounds by live cell imaging. In the absence of BRD-1 we observed diffuse fluorescence within the nucleoplasm from proliferative zone to mid-pachytene, with no evidence of tracks (Fig 3B). In late pachytene, some regions of more intense GFP::BRC-1 fluorescence were observed; however, in diplotene and diakinesis only a diffuse nuceloplasmic signal was detected, with no concentrated regions of GFP::BRC-1. This indicates that BRD-1 is required for the correct localization of BRC-1 in meiotic cells. In contrast, BRD-1::GFP fluorescence in *brc-1* mutants appeared similar to wild type (Fig 3B). Analysis of steady state protein levels by immunoblot revealed that BRC-1 and BRD-1 are relatively stable in the absence of the other partner (*brc-1* = 86% of BRD-1::GFP levels and *brd-1* = 74% of GFP::BRC-1 levels compared to wild-type extracts; Fig 3C). Thus, these results suggest that BRD-1 is uniquely required for localization of the complex to chromatin and the SC.

### Impairment of either meiotic recombination or synaptonemal complex formation alters GFP::BRC-1 localization

To provide insight into the relationship between BRC-1/BRD-1 and the progression of meiotic recombination, we monitored the localization of GFP::BRC-1 in mutants that impair different steps of meiotic recombination: *spo-11* mutants are unable to form meiotic DSBs (32, 33), *rad-51* mutants are blocked prior to strand invasion (34-36), and *msh-5* mutants fail to form crossovers (37, 38). In *spo-11* mutants, we observed many fewer GFP::BRC-1 foci in transition zone and early pachytene compared to WT (WT = 78.87±3.84% vs. *spo-11* = 23.83±2.12% of nuclei had one or more foci; n=4 germ lines; p = 0.0002). At mid-pachytene GFP::BRC-1 was observed in tracks in the *spo-11* mutant similar to wild type, as synapsis occurs in the absence of genetic recombination in *C. elegans* (32) (Fig 4). In late pachytene, GFP::BRC-1 fluorescence did not concentrate on a portion of each chromosome pair nor retract to the short arm of the bivalent as in wild type, consistent with these events being dependent on crossover formation. However, in 20.23±1.78% of nuclei (n=4 germ lines) there was enrichment of GFP::BRC-1 on a single chromosome, as has been previously observed for synapsis markers, including the phosphorylated form of SYP-4 (39, 40), and likely represents *spo-11*-independent lesions capable of recruiting meiotic DNA repair components and altering SC properties. Consistent with this, we observed co-localization of the concentrated GFP::BRC-1 and phospho-SYP-4 on the occasional chromosome track (S2 Fig). As expected, BRD-1::GFP was observed in a similar pattern to GFP::BRC-1 in *spo-11* mutants throughout meiotic prophase (S2 Fig).

**Fig 4.**
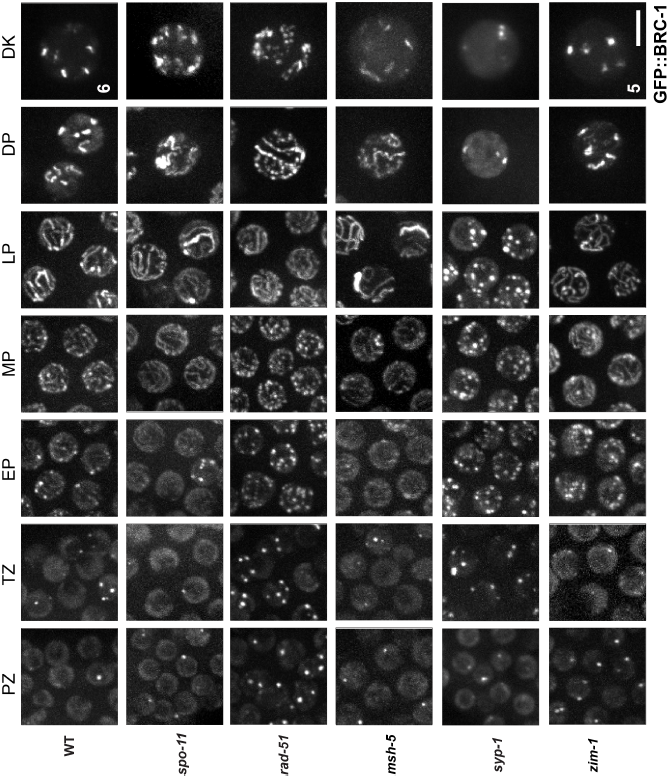
GFP::BRC-1 localization is perturbed when either meiotic recombination or chromosome synapsis is impaired. High-magnification images of live *C. elegans* expressing GFP::BRC-1 from the indicated genetic backgrounds and gonad region (PZ = Proliferative Zone, TZ = Transition Zone, EP = Early Pachytene, MP = Mid Pachytene, LP = Late Pachytene, DP = Diplotene, DK = Diakinesis). In wild type GFP::BRC-1 localizes to chromatin and in a small number of foci in the proliferative and transition zone. When synapsis is initiated in early and mid pachytene, GFP::BRC-1 begins to localize in long tracks corresponding to the synaptonemal complex. In late pachytene, GFP::BRC-1 becomes condensed in shorter regions along the length of synapsed chromosomes, demarcating what will become the short arms of the six bivalent chromosomes in diakinesis. This localization pattern is perturbed when synapsis and crossover formation are disrupted. Scale bar = 5 μm.

Following DSB formation and processing, RAD-51 is loaded onto resected single-stranded DNA and facilitates strand exchange (36). GFP::BRC-1 localization was significantly impaired in the *rad-51* mutant (Fig 4). More nuclei contained GFP::BRC-1 foci throughout the germ line. In transition zone/early pachytene 95.87±0.095% of nuclei had foci, compared to 78.87±3.84% in WT (n=3 *rad-51* germ lines; p = 0.0127). These foci presumably represent resected DSBs that fail to undergo strand invasion in the absence of RAD-51. Track-like structures were not observed until late pachytene in the absence of RAD-51. The punctate nature of GFP::BRC-1 was particularly pronounced in diplotene and diakinesis, with no clear concentration to six regions, consistent with the absence of crossovers in the *rad-51* mutant.

In *msh-5* mutants, GFP::BRC-1 appeared similar to wild type from the proliferative zone to mid pachytene, localizing in the nucleoplasm and concentrating in foci before converging on tracks (Fig 4). Similar to *spo-11*, 26.27±2.25% of *msh-5* late pachytene nuclei (n=4 germ lines) contained concentrated GFP::BRC-1 on one or occasionally two tracks. This contrasts with synapsis markers, which are not enriched on any chromosome in the absence of factors required for crossover formation (39, 40). Therefore, while blocking crossover formation impairs BRC-1 localization, it can still concentrate on a subset of chromosomes. Taken together, our data suggest that GFP::BRC-1 localizes to the SC and its concentration and retraction to the short arm of the bivalent is dependent on processing of meiotic DSBs.

We also examined localization of GFP::BRC-1 when synapsis is blocked by mutation of a component of the central region of the SC, *syp-1* (29). GFP::BRC-1 is nucleoplasmic in the absence of synapsis and concentrates in foci in every nucleus throughout meiotic prophase (n=3 germ lines; Fig 4). However, it never becomes associated with tracks, nor concentrates or retracts on chromosomes. Thus, GFP::BRC-1 localization to tracks is dependent on SC formation.

To examine localization under conditions where a subset of chromosomes fail to synapse and recombine, we monitored GFP::BRC-1 localization in the *zim-1* mutant, in which chromosomes *II* and *III* cannot synapse (41). In early pachytene, GFP::BRC-1 was observed in many foci in the *zim-1* mutant, similar to the *syp-1* mutant (Fig 4). However, as meiosis progressed, GFP::BRC-1 was observed on tracks that condensed to the short arm of the bivalent on multiple chromosomes. Many times we observed more than four stretches of GFP::BRC-1 fluorescence at diplotene/diakinesis (Fig 4), suggesting that there are more than four chiasmata in the *zim-1* mutant. We address the altered number of chiasmata in the *zim-1* mutant below.

### BRC-1/BRD-1 is important when chromosome synapsis and crossovers are perturbed and alters RAD-51 patterns

Given the association of GFP::BRC-1 and BRD-1::GFP with the SC and its alteration when synapsis is perturbed (Fig 4), we next examined the functional consequence of removing BRC-1/BRD-1 in the *zim-1* mutant. Inactivation of BRC-1 or BRD-1 in *zim-1* led to enhanced embryonic lethality compared to the single mutants, suggesting that BRC-1/BRD-1 plays a role in promoting viability when chromosomes are unable to synapse and recombine (p<0.0001; Fig 5A).

**Fig 5.**
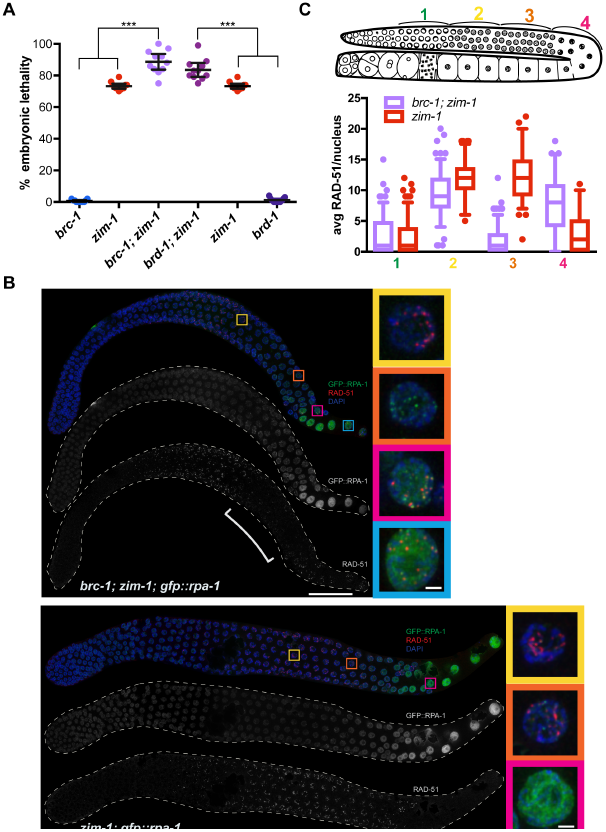
*brc-1* and *brd-1* enhance embryonic lethality and alter the pattern of RAD-51 in the *zim-1* mutant. A) Embryonic lethality in indicated mutants; 95% Confidence Intervals are shown. The genetic interaction between *brc-1/brd-1* and *zim-1* is significant by a one-way ANOVA (*** = p<0.0001). B) Dissected germ lines from *brc-1; zim-1; gfp::rpa-1* and *zim-1; gfp::rpa-1* worms stained with anti-RAD-51 (red) and GFP::RPA-1 fluorescence (green), counterstained with DAPI (blue). Scale bar = 20 μm. Insets show selected nuclei from different regions of the germ line; bracket indicates RAD-51 “dark zone”. Scale bar = 1 μm. C) Schematic of germ line indicating zones for analysis of RAD-51 foci. Box whisker plots show average number of RAD-51 foci per nucleus in the different zones. Horizontal line of each box indicates the median, the top and bottom of the box indicates medians of upper and lower quartiles, lines extending above and below boxes indicate standard deviation and individual data points are outliers from 5-95%. Statistical comparisons by Mann-Whitney of *brc-1;zim-1* versus *zim-1* in the different regions of the germ line: 1, p=0.239; 2, p<0.0001; 3, p<0.0001; 4, p<0.0001.Numbers of nuclei scored in each zone for *brc-1; zim-1*: 1 = 177; 2 = 138; 3 = 161; 4 = 61; *zim-1*: 1 = 159; 2 = 88; 3 = 103; 4 = 78.

To determine the nature of the enhanced embryonic lethality of *zim-1* in the absence of *brc-1/brd-1*, we monitored RAD-51 assembly/disassembly in the spatiotemporal organization of the germ line. Previous analyses revealed that *brc-1* mutant hermaphrodites have elevated RAD-51 foci in late pachytene, suggesting that repair of a subset of meiotic DSBs is delayed in the absence of BRC-1 (9), which we also observed in both the *brc-1* and *brd-1* mutants (S3 Fig).Further, blocking synapsis on some or all chromosomes results in elevated RAD-51 levels genome wide (28, 42), which we also observed in the *zim-1* mutant (Fig 5B, C). Surprisingly, removal of BRC-1 or BRD-1 in *zim-1* led to fewer RAD-51 at mid-late pachytene: RAD-51 foci appeared at similar levels compared to the *zim-1* single mutant early in meiotic prophase, but in the latter half of pachytene much of RAD-51 was no longer detected on chromosomes. High levels of RAD-51 were observed again at the gonad bend, as nuclei exited pachytene and entered diplotene (Fig 5B, C). Similar patterns were observed for *brd-1; zim-1* as well as when *brc-1* was removed in other mutants that perturb synapsis (S3 Fig). These results suggest that when crossover formation is perturbed by blocking synapsis, BRC-1/BRD-1 plays a role in DSB formation, DNA end resection, RAD-51 loading, and/or stabilization of the RAD-51 filament in mid-late pachytene.

To differentiate between these possibilities for BRC-1/BRD-1 function, we first analyzed the pattern of the single-stranded binding protein RPA-1 [GFP::RPA-1; (43)]. RPA-1 binds resected ends prior to RAD-51 loading (44) and is also associated with recombination events at a post-strand-exchange step, which can be observed in chromosome spreads (45). In the *brc-1; zim-1* germ line we observed an inverse pattern between RAD-51 and RPA-1 at mid-late pachytene: GFP::RPA-1 foci were prevalent in the region where RAD-51 foci were reduced (Fig 5B). In the *zim-1* single mutant, fewer GFP::RPA-1 foci were observed at this stage, while RAD-51 remained prevalent. We also observed few RPA-1 foci at mid-late pachytene in wild type or the *brc-1* single mutant whole mount gonads (S2 Fig). These results suggest that BRC-1 is not required for break formation *per se* in this region of the germ line, as we observed an increase in GFP::RPA-1 foci, not a decrease as would be expected if BRC-1/BRD-1 is required for DSB formation. Additionally, this result argues against a role for BRC-1/BRD-1 in promoting resection as RPA-1 loads on exposed single stranded DNA (44). Thus, at mid to late pachytene BRC-1/BRD-1 either facilitates the assembly of RAD-51 on new breaks, and/or stabilizes the RAD-51 filament.

### BRC-1/BRD-1 stabilizes the RAD-51 filament when crossover formation is perturbed

The lack of RAD-51 in mid to late pachytene in *brc-1; zim-1* is reminiscent of the RAD-51 “dark zone” observed in the *spo-11; rad-50* mutant following exposure to IR; this likely reflects a requirement for RAD-50 in loading RAD-51 at resected DSBs on meiotic chromosomes (46). However, the distal boundary of the dark zone in the *brc-1; zim-1* double mutant is distinct from the *rad-50* mutant: the dark zone in *rad-50* extends from meiotic entry to late pachytene (46), while in *brc-1; zim-1,* reduction in RAD-51 is limited to mid-late pachytene (Fig 5B, C), suggesting that the nature of the dark zone is different in these mutant situations. If BRC-1/BRD-1 was required for loading RAD-51 on breaks in mid-late pachytene, then a time course analysis would reveal a diminution of the dark zone by twelve hours following IR exposure, as was observed for *spo-11; rad-50* mutants (Fig 6A, loading defect on left) (46). On the other hand, if BRC-1 was important for protecting RAD-51 from disassembly, then the dark zone should be maintained throughout the time course as RAD-51 would be disassembled as nuclei with pre-installed RAD-51 move through the mid-late pachytene region of the germ line (Fig 6A, stabilization defect on right). To examine this, we exposed *spo-11* and *brc-1*; *spo-11* mutants depleted for SYP-2, where all chromosomes fail to synapse and therefore do not form crossovers, to 10 Gys of IR and examined RAD-51 at 1, 4, 8, and 12 hours following IR exposure. The *spo-11* background eliminates meiotic DSB formation (32), which remains active under conditions where crossovers have not formed on all chromosomes (*e.g*., *syp-2*) (47, 48). Thus, under this scenario breaks are induced uniformly in the germ line at a single point in time and as nuclei move through the germ line, no new breaks are formed. At 1, 4, 8, and 12 hours following IR, the dark zone was maintained in the absence of BRC-1 (Fig 6B). This result is consistent with the hypothesis that BRC-1/BRD-1 stabilizes the RAD-51 filament rather than facilitates loading of RAD-51 on new DSBs at mid-late pachytene.

**Fig 6.**
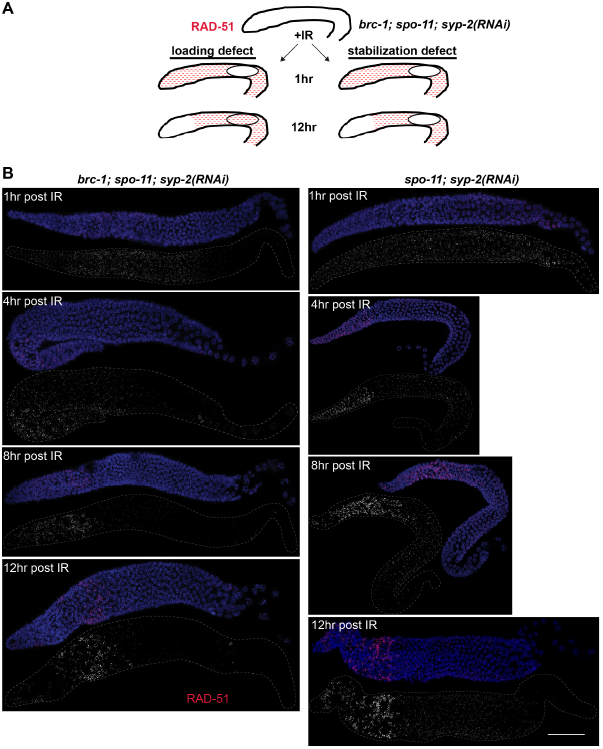
BRC-1 promotes the stability of RAD-51 when crossover formation is impaired. A) Schematic of potential outcomes of IR-induced RAD-51 (red) over time in *brc-1; spo-11; syp-2*(RNAi): a defect in RAD-51 loading would result in loss of the “dark zone” (circle) by 12 hrs (left), while a defect in RAD-51 stabilization would manifest in the maintenance of the dark zone over 12 hrs (right). B) Projections of whole germ lines at indicated times after IR treatment in *brc-1; spo-11; syp-2*(RNAi) (left) and *spo-11; syp-2*(RNAi) (right). Scale bar = 30 μm.

### BRC-1/BRD-1 alters recombination outcomes

A subset of RAD-51 strand invasions are processed into crossovers, which are marked by the cyclin related protein, CNTD1/COSA-1 (31, 49). Given the reduction in RAD-51 in mid-late pachytene in *brc-1; zim-1* mutant hermaphrodites, we next analyzed crossover precursor formation in the various mutants. In *C. elegans*, each of the six chromosome pairs has a single crossover; consequently, there are six COSA-1 foci in hermaphrodite germ cells at late pachytene (31) (5.996±0.004; Fig 7A). We also observed six COSA-1 foci in late pachytene nuclei in the *brc-1* and *brd-1* mutants (6.009 ± 0.003 and 5.979 ± 0.015, respectively; Fig 7A), indicating that breaks are efficiently processed into crossovers in the absence of BRC-1/BRD-1 in an otherwise wild-type worm. This is consistent with the presence of six bivalents at diakinesis and the low embryonic lethality of *brc-1* and *brd-1* (9, 10) (S1 Fig). In *zim-1* mutants we expected to observe four COSA-1 foci per nucleus marking the four paired chromosomes, but not the unpaired chromosome *II*s and *III*s. Contrary to our expectations, *zim-1* had 6.117±0.119 COSA-1 foci (chi square from expected, p<0.005), with a very broad distribution ranging from 2 to 9 foci; such a wide distribution is never observed in wild type (31) (Fig 7A; S4 Fig). Inactivation of BRC-1 in *zim-1* mutants reduced the number of GFP::COSA-1 foci to 4.831±0.068, closer to expectations although still significantly different than expected (chi square p<0.005), and the distribution remained broad (p <0.0001; Fig 7A; S4 Fig). Very similar results were obtained for *brd-1; zim-1* mutants (Fig 7A). These results suggest that when crossovers are unable to form between some homologs, additional COSA-1-marked crossover precursors are generated, and some of these are dependent on BRC-1/BRD-1.

**Fig 7.**
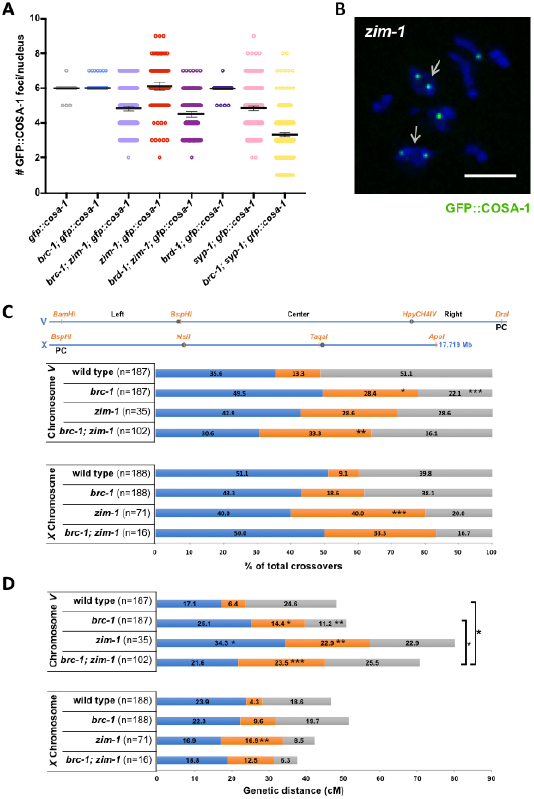
BRC-1/BRD-1 alters the crossover landscape. A) Number of COSA-1 foci in mid-late pachytene in indicated mutants. Number of nuclei scored: *gfp::cosa-1* = 458, *brc-1; gfp::cosa-1*= 815, *brc-1; zim-1; gfp::cosa-1* = 255, *zim-1; gfp::cosa-1* = 120, *brd-1; zim-1*; *gfp::cosa-1* = 164, *brd-1; gfp::cosa-1* = 145, *syp-1; gfp::cosa-1* = 292, *syp-1; brc-1; gfp::cosa-1* = 487. B) Diplotene *zim-1; gfp::cosa-1* nucleus showing ring chromosomes (arrows) and GFP::COSA-1. Scale bar = 2 μm. C) SNP markers and distribution of crossovers on chromosome *V* and the *X* chromosome in wild type, *brc-1, zim-1* and *brc-1; zim-1* mutants. D) The genetic map is expanded in *zim-1*, and this is partially dependent on BRC-1.

The higher than expected numbers of COSA-1 foci observed in *zim-1* mutants could reflect recombination intermediates that do not go on to form chiasmata (i.e., non-crossovers or inter-sister crossovers), or could be *bona fide* inter-homolog crossovers, such that some chromosomes have more than one chiasma. To provide insight into the nature of the extra COSA-1 foci, we analyzed COSA-1 in *syp-1* mutants, where no chiasmata can form as all chromosomes fail to synapse, and found that there were on average 4.85±0.07 COSA-1 foci at late pachytene (Fig 7A; S4 Fig). These results suggest that under conditions of meiotic dysfunction when chiasmata cannot form because chromosomes are unable to pair/synapse, COSA-1 is recruited to recombination events that become processed into non-crossovers and/or inter-sister crossovers. As with *zim-1* mutants, inactivation of BRC-1 in the *syp-1* mutant background led to fewer COSA-1 foci (*brc-1; syp-1*: 3.32±0.052 vs. *syp-1*: 4.846±0.069, p<0.0001) (Fig 7A; S4 Fig), consistent with BRC-1 promoting recombination events under conditions where chiasma formation is blocked.

To determine whether the extra COSA-1 foci on synapsed chromosomes could form chiasmata, we examined *zim-1* and *brc-1; zim-1* diplotene/diakinesis nuclei, where chromosomes are individualized and cross-shaped structures indicative of crossovers between homologs can be observed. Consistent with the formation of extra chiasmata in the *zim-1* mutant background, we observed 52% of diplotene/diakinesis nuclei (n = 52) containing at least one ring-shaped structure, and six had two ring-shaped structures; the simplest interpretation is that there was a chiasma on each end of the chromosome pair (arrow; Fig 7B). This was reduced to 21% of diplotene/diakinesis nuclei (n = 43) containing ring-shaped chromosomes in the *brc-1; zim-1* double mutant (*zim-1* vs. *brc-1; zim-1*, p=0.0028 Mann-Whitney). These results suggest that BRC-1 promotes chiasma formation under conditions where some chromosomes are unable to interact with their partner.

To genetically examine crossovers, we monitored linkage between SNP makers on chromosomes *V* and *X* in Bristol/Hawaiian hybrid strains to assess both crossover numbers and distribution. While inactivation of *brc-1* had no effect on crossover numbers on either chromosome *V* or *X*, we observed an altered distribution of crossovers on chromosome *V* in the *brc-1* mutant (Fig 7C; S1 Table). In *C. elegans*, crossovers are enriched on the arms, presumably due to the necessity to reorganize meiotic chromosomes into long and short arms for regulated sister chromatid release and segregation (30, 50-52). In the *brc-1* mutant we observed a more even distribution of crossovers, with more crossovers in the center and fewer on the right arm (Fig 7C; S1 Table). On the other hand, crossover distribution was not significantly different on the *X* chromosome, which has an altered crossover landscape (53, 54), in the absence of BRC-1.

To determine whether the extra COSA-1 foci and ring-shaped chromosomes observed in *zim-1* result in genetic exchange, we monitored linkage between SNP markers in the *zim-1* and *brc-1; zim-1* mutants. Analysis of the *zim-1* mutant revealed an increase in the recombination map on chromosome *V*, and multiple double crossovers were observed (Fig 7D; S1 Table). Extra crossovers were also observed on autosomes in worms unable to pair and synapse *X* chromosomes (42). Inactivation of BRC-1 in the *zim-1* background resulted in fewer crossovers (Fig 7D; S1 Table). We observed no significant differences on the *X* chromosome, although this may be a consequence of the small numbers analyzed.

*C. elegans* has very strong interference, which is the phenonmon that a crossover at one position on a chromosome decreases the probability of formation of a crossover nearby, resulting in a single crossover per chromosome (52). Given the increase in double crossovers observed in the *zim-1* mutant on chromosome *V*, we calculated the interference ratio. While wild type and *brc-1* had absolute intereference of 1, as no double crossovers were observed, the *zim-1* muntant no longer displayed interference (Table 1). Inactivation of BRC-1 in the *zim-1* mutant resulted in partial suppression of the interference defect (Table 1). Together, these results suggest that BRC-1 counteracts interference to promote crossover formation under conditions of meiotic dysfunction.

**Table 1.** *brc-1* partially suppresses interference in the *zim-1* mutant.

| WT (V) | expected DCO | observed DCO | c.o.c. | interference |
| --- | --- | --- | --- | --- |
| LC | 0.01 | 0.00 | 0.00 | 1.00 |
| CR | 0.02 | 0.00 | 0.00 | 1.00 |
| LR | 0.04 | 0.00 | 0.00 | 1.00 |
| *brc-1* (V) | expected DCO | observed DCO | c.o.c. | interference |
| LC | 0.04 | 0.00 | 0.00 | 1.00 |
| CR | 0.02 | 0.00 | 0.00 | 1.00 |
| LR | 0.03 | 0.00 | 0.00 | 1.00 |
| *zim-1*(V) | expected DCO | observed DCO | c.o.c. | interference |
| LC | 0.08 | 0.09 | 1.09 | -0.09 |
| CR | 0.05 | 0.03 | 0.55 | 0.45 |
| LR | 0.08 | 0.11 | 1.46 | -0.46 |
| *brc-1;zim-1*(V) | expected DCO | observed DCO | c.o.c. | interference |
| LC | 0.05 | 0.03 | 0.58 | 0.42 |
| CR | 0.06 | 0.03 | 0.49 | 0.51 |
| LR | 0.05 | 0.03 | 0.53 | 0.47 |

LC = left-center interval; CR = center-right interval; LR = left-right interval. DCO: double crossover; expected DCO: (crossover frequency at interval "A") x (crossover frequency at interval "B"). c.o.c. (coefficient of coincidence) = actual DCO frequency/ expected DCO frequency; Interference = 1- c.o.c.

## Discussion

Here we show that *C. elegans* BRC-1 and BRD-1 orthologs localize to the SC and promote homologous recombination, particularly when meiosis is perturbed. These results suggest that BRC-1/BRD-1 plays a critical checkpoint role in monitoring and modulating DSB repair in the context of the specialized meiotic chromosome structure.

### BRC-1/BRD-1 localizes to synapsed chromosomes and its retraction to the short arm of the bivalent precedes reorganization of the SC

In mouse spermatocytes BRCA1 is associated with RAD51 and enriched on asynapsed regions of meiotic chromosomes, including the X-Y sex body (18, 19). Here we show that *C. elegans* BRC-1 and BRD-1 partially associate with RAD-51 in early meiotic prophase, but become enriched on synapsed chromosomes as meiosis progresses, co-localizing with SYP-1, a SC central region component (Fig 2). The enrichment of mammalian BRCA1 on asynapsed chromosomes versus on synapsed chromosomes in *C. elegans* most likely reflects alteration in the relationship between meiotic recombination and SC formation in these organisms. Meiotic chromosomes can pair and synapse in the absence of meiotic recombination in *C. elegans* (32), while these events are interdependent in mammals (15, 16). The HORMAD axial components also show differences in chromosome association in mice and worms: in mice, HORMAD1 and 2 are enriched on asynapsed chromosomes (55, 56), while *C. elegans* HORMADS, HIM-3, HTP-1/2, and HTP-3, remain associated with synapsed chromosomes (57-60). However, the function of HORMADs in preventing inter-sister recombination and in checkpoint signaling appears to be similar in these different organisms (61-66). Thus, the association of BRC-1/BRD-1 to the SC in *C. elegans* is likely a consequence of the inter-relationship between SC formation and meiotic recombination in this organism and not in different functions for this complex in worm versus mammalian meiosis. As women carrying BRCA1 mutations with no indication of cancer have decreased ovarian reserve, accelerated primordial follicle loss, and oocyte DNA damage (67), it is possible that the BRCA1/BARD1 complex plays similar roles in human and worm meiosis.

Another difference between *C. elegans* and mammals is the nature of the kinetochore. *C. elegans* chromosomes are holocentric while in many organisms, including yeast and mice, chromosomes are monocentric. Holocentricity dictates that a single off-centered crossover is formed on each homolog pair to define the long and short arms necessary to ensure regulated sister chromatid cohesion release at meiosis I and II (50-52, 68). Interestingly, BRC-1/BRD-1 becomes restricted to the short arm of the bivalent, as defined by the crossover site, and this precedes SC reorganization. While the absence of BRC-1/BRD-1 alone does not overtly affect crossover formation (Fig 7), it does alter the distribution of crossover events along chromosome *V* such that more events occur in the middle of the chromosome. The change in crossover distribution in *brc-1* mutants may contribute to the increased nondisjunction observed in the absence of the BRC-1/BRD-1 complex.

We show that the concentration of BRC-1/BRD-1 to a portion of each chromosome track in late pachytene is dependent on meiotic DSB processing (Fig 4). This is similar to what has recently been shown for SC components, such as the phosphorylated form of the central region component SYP-4 (39, 40). While RAD-51 is absolutely required for the concentration and retraction of BRC-1 and phospho-SYP-4, in *spo-11* mutants occasional chromosomes show BRC-1/BRD-1 and phospho-SYP-4 concentration, although no retraction (39) (S2A Fig). This most likely reflects the ability of *spo-11*-independent lesions to recruit meiotic DNA repair components (39, 40). However, in contrast to phospho-SYP-4, BRC-1 also concentrates on occasional chromosomes in the absence of crossover factors. This suggests that BRC-1 can respond to other repair pathways besides the formation of inter-homolog crossovers. One possibility is that when inter-homolog crossover formation is blocked, DSBs are repaired through site-specific nucleases (69-71), a subset of which leads to the concentration of BRC-1/BRD-1 on chromosomes.

### BRC-1 and BRD-1 are not identical and may contribute different functions to the complex

BRCA1 forms a potent E3 ubiquitin ligase only in complex with its partner BARD1 (2, 3). Biochemical and structural studies have defined the RING domains and associated helices of these proteins as critical for catalytic activity and BRCA1-BARD1 interaction (72). However, while the BRCA1/BARD1 heterodimer exhibits substantially greater E3 ligase activity *in vitro* than BRCA1 alone, only the BRCA1 RING domain interacts with the E2 for ubiquitin transfer, suggesting that BRCA1 is the critical subunit for E3 ubiquitin ligase activity (3, 73). Structure-function analysis of the BARD1 RING domain suggests that BARD1 may serve to attenuate BRCA1 E3 ligase activity (74), reinforcing the premise that these proteins do not serve the same function within the complex. We found that while *brc-1* and *brd-1* mutants have very similar meiotic phenotypes (Figs S1, 5, 6, S3, 7), and the localization of BRC-1 is dependent on BRD-1, BRD-1 localization to meiotic chromosomes is independent of BRC-1 function (Fig 3C). This is in contrast to a previous report where BRC-1 was shown to be required for DNA damage-induced BRD-1 foci formation (27) and the findings in the accompanying paper by Janisiw et al. Both of these studies used antibodies directed against BRD-1, which requires fixation that can alter antigen-antibody interaction, while we monitored functional fluorescent fusion proteins in live worms. Alternatively, the differences we observed could be a consequence of appending GFP to BRD-1, which can result in dimerization and stabilization of the fusion protein (75).Nonetheless, these results suggest that BRD-1 has different properties than BRC-1 with respect to association with meiotic chromosomes.

Similar to the mammalian proteins, both BRC-1 and BRD-1 contain long linker and phosphoprotein binding BRCA1 C-terminal (BRCT) domains in addition to the N-terminal RING domains that confer E3 ligase activity. BRCT domains are phosphorylation-dependent interacting modules that have been implicated in tumor suppressor activity (76). Interestingly, only BRD-1 contains Ankyrin (ANK) repeat interaction domains. Recent structural and functional analyses of the ANK domain in TONSL-MMS22L, a complex involved in homologous recombination, revealed that the ANK domain interacts with histone H4 tails (77). The BARD1 ANK domains have a very similar fold (77), suggesting that BARD1 ANK domains may be important for association with chromatin. Future work will determine whether the ANK domains in BRD-1 mediate the association of this complex to meiotic chromosomes and may uncover other BRD-1-specific functions.

### BRC-1/BRD-1 function in meiotic recombination

It has long been appreciated that BRCA1/BARD1 mediates its tumor suppressor activity at least in part through regulating homologous recombination (6). Given the importance of homologous recombination in promoting chiasma formation during meiosis, it is not surprising that removing BRC-1/BRD-1 impinges on meiotic recombination. BRCA1/BARD1 associates with the key recombinase RAD51 in both mammals and *C. elegans* (19, 27, 78). BRCA1 has also been shown to be required for the assembly of DNA damage induced RAD51 foci in chromatin (79), and this has been interpreted as a requirement for BRCA1 in RAD51 filament assembly. However, recent biochemical analyses using purified proteins found that BRCA1 is not required for RAD51 assembly on RPA coated single stranded DNA and instead promotes DNA strand invasion (78). Further, a BARD1 mutant that cannot interact with RAD51 does not promote DNA strand invasion, and also does not form foci *in vivo.* Thus, it is likely that BRCA1/BARD1 is not required for RAD51 filament assembly *per se*. Our IR time course analysis of *C. elegans spo-11; brc-1* mutants is consistent with a function for this complex in stabilizing the RAD-51 filament. It is possible that similar to the mammalian complex, BRC-1/BRD-1 promotes RAD-51 strand invasion; however, *in vivo* the RAD-51 filament may be subject to disassembly by other proteins in the absence of BRC-1/BRD-1, which would not be recapitulated *in vitro*. One such protein is the FANCJ/DOG-1 helicase, which interacts with BRCA1 (80), and can disassemble RAD51 on ssDNA *in vitro* (81). It is also likely that BRCA1/BARD1 plays multiple roles during homologous recombination and interacts with, and coordinates the activity, of many proteins, including RAD51, and these interactions are modulated under different conditions, including DNA damage, meiosis, meiotic dysfunction, as well as at different stages of the cell cycle. Consistent with this, Janisiw et al. found that BRC-1 associates with the pro-crossover factor MSH-5.

### BRCA1/BARD1 serves a checkpoint function in meiosis

*brc-1* and *brd-1* mutants have very subtle defects in an otherwise wild-type meiosis. This includes low levels of chromosome nondisjunction (10) (S1 Fig), a delay in repair of a subset of DSBs through the inter-sister pathway (9), and elevated heterologous recombination (12). However, the effect of removing BRC-1/BRD-1 when meiosis is perturbed in mutants that impair chromosome pairing, synapsis and crossover recombination leads to enhanced meiotic dysfunction, including elevated embryonic lethality (Fig 5), impaired RAD-51 stability (Fig 6), and alteration of COSA-1 and the crossover landscape (Fig 7), suggesting that BRC-1/BRD-1 functions is critical when meiosis is perturbed.

In both *C. elegans* and *Drosophila melanogaster*, preventing crossover formation on a subset of chromosomes leads to additional events on other chromosomes, and is referred to as the interchromosomal effect (42, 82-84). There is also evidence in humans that Robertsonian translocations elicit the interchromosomal effect (85). Our analysis of the *zim-1* mutant, where chromosomes *II* and *III* fail to recombine, revealed elevated COSA-1 foci genome wide and an increase in genetic crossovers on chromosome *V* (Fig 7), consistent with the interchromosomal effect. These results also reveal that when meiosis is perturbed as in *syp-1,* and perhaps *zim-1* mutants, COSA-1 can mark events that don’t ultimately become crossovers. Interestingly, removal of BRC-1 in the *zim-1* mutant decreased both the number of COSA-1 foci and crossovers, suggesting that BRC-1 plays a role in the interchromosomal effect. One consequence of the interchromosomal effect is that the potent crossover interference observed in *C. elegans*, which limits crossovers to one per chromosome (52), appears to be alleviated when some chromosomes cannot form crossovers. The reduction of crossover numbers in the *brc-1* mutant suggests that BRC-1/BRD-1 functions as an anti-interference factor under conditions of meiotic dysfunction. As crossover interference is mediated by meiotic chromosome structure (86), it is likely that SC-associated BRC-1/BRD-1 counteracts interference when meiosis is perturbed.

In conclusion, our results indicate that BRC-1/BRD-1 serves a critical role in monitoring the progression of meiotic recombination in the context of the SC when meiosis cannot proceed normally, suggesting that BRC-1/BRD-1 serves a checkpoint function. When crossover formation is blocked, BRC-1/BRD-1 stabilizes the RAD-51 filament and promotes processing of DSBs by homologous recombination, some of which go on to form additional crossovers on synapsed chromosomes. In this context, BRC-1/BRD-1 joins a growing list of proteins that monitor meiotic recombination to promote accurate chromosome segregation (47, 48, 87-90). Future work will examine the relationship between BRC-1/BRD-1 and other meiotic checkpoint pathways and identify substrates of BRC-1/BRD-1-ubiquitination to understand how this complex modulates recombination under conditions when meiosis is perturbed.

## Materials and Methods

### Generation of GFP::BRC-1, RFP::BRC-1 and BRD-1::GFP

Fusions were generated as described in Dickenson (20) and backcrossed a minimum of three times. Sequence information is available upon request.

### Genetics

*C. elegans* var. Bristol (N2), was used as the wild-type strain. Other strains used in this study are listed in **S2 Table**. Some nematode strains were provided by the Caenorhabditis Genetics Center, which is funded by the National Institutes of Health National Center for Research Resources (NIH NCRR). Strains were maintained at 20°C.

### Embryonic lethality and production of male progeny

Embryonic lethality in the absence or presence of 5mM hydroxyurea (16 hrs), or 75 Grays (Gys) of γ-irradiation (IR) from a ^137^Cs source, was determined over 3 days by counting eggs and hatched larvae 24 hrs after removing the hermaphrodite and calculating percent as eggs/eggs + larvae; male progeny was assessed 48 hrs after removing the hermaphrodite. A minimum of 10 worms were scored for each condition.

### Cytological Analysis

Immunostaining of germ lines was performed as described (91) except slides were incubated in 95% ethanol instead of 100% methanol for direct GFP fluorescence of GFP::BRC-1, BRD-1::GFP, GFP::RPA-1, and GFP::COSA-1. Staining with antibodies against phospho-SYP-4 was as described (39). The following primary antibodies were used at the indicated dilutions: rabbit anti-RAD-51 (1:10,000; Catalog #29480002) and rabbit anti-GFP (1:500; NB600-308) (Novus Biologicals, Littleton, CO), mouse anti-GFP (1:500) (Millipore, Temecula, CA), mouse anti-α-tubulin (1:500; Sigma-Aldrich; T9026), anti-SYP-1 (1:200) (generously provided by Anne Villeneuve); rabbit anti-phospho-SYP-4 (1:100; (39)), guinea pig anti-HTP-3 (1:500) (generously provided by Abby Dernburg). Secondary antibodies Alexa Fluor 594 donkey anti-rabbit IgG and Alexa Fluor 488 goat anti-guinea pig IgG from Life Technologies were used at 1:500 dilutions. DAPI (2μg/ml; Sigma-Aldrich) was used to counterstain DNA.

Collection of fixed images was performed using an API Delta Vision deconvolution microscope or a Nikon TiE inverted microscope stand equipped with an 60x, NA 1.49 objective lens, Andor Clara interline camera, motorized and encoded stage, and appropriate filters for epi-fluorescence. Z stacks (0.2 μm) were collected from the entire gonad. A minimum of three germ lines was examined for each condition. Images were deconvolved using Applied Precision SoftWoRx or Nikon NIS Elements Offline batch deconvolution software employing either “Automatic3D” or “Richardson-Lucy” deconvolution modes and subsequently processed and analyzed using Fiji (ImageJ) (Wayne Rasband, NIH).

RAD-51 foci were quantified in a minimum of three germ lines of age-matched hermaphrodites (18-24 hr post-L4). As *zim-1* mutants have an extended transition zone (41), we divided germlines into four equal zones from the beginning of the transition zone (leptotene/zygotene), as counted from the first row with three or more crescent-shaped nuclei, through diplotene (Fig 5C). The number of foci per nucleus was scored for each region.

To assess formation of RAD-51 foci following IR treatment, 18-24 hrs post-L4 worms were exposed to 75 Grays (Gys) of IR; gonads were dissected and fixed for immunofluorescence as above 2 hrs post IR. Time course analysis was performed by exposing worms to 10 Gys and dissecting 1, 4, 8, and 12 hrs following IR treatment.

For live cell imaging, 18-24 hours post L4 hermaphrodites were anesthetized in 1mM tetramisole (Sigma-Aldrich) and immobilized between a coverslip and an 1.5% agarose pad on a glass slide. Z-stacks (0.33 μm) were captured on a spinning-disk module of an inverted objective fluorescence microscope [Marianas spinning-disk confocal (SDC) real-time 3D Confocal-TIRF (total internal reflection) microscope; Intelligent Imaging Innovations] with a 100×, 1.46 numerical aperture objective, and a Photometrics QuantiEM electron multiplying charge-coupled device (EMCCD) camera. Z-projections of approximately 20-30 z-slices were generated, cropped, and adjusted for brightness in ImageJ.

### Immunoblot analysis

Whole worm lysates were generated from indicated worms; unmated *fog-2(q71)* worms were used to eliminate embryos. Lysates were resolved on 4-15% SDS-PAGE gradient gels (Bio-RAD) and transferred to Millipore Immobilon-P PVDF membranes. Membranes were blocked with 5% BSA and probed with indicated antibodies, including anti-α-tubulin as loading control, followed by IRDye680LT-and IRDye800-conjugated anti-rabbit and anti-mouse IgG secondary antibodies obtained from LI-COR Bioscience (Lincoln, NE).Immunoblots were repeated a minimum of three times, and signal was quantified using ImageJ and normalized with the α-tubulin signal.

### RNA-mediated interference analysis

RNA-mediated interference (RNAi) was performed at 20°C, using the feeding method (92). Cultures were plated onto NGM plates containing 25μg/ml carbenicillin and 1 mM IPTG and were used within 2 weeks.

### Meiotic mapping

Meiotic crossover frequencies and distribution were assayed utilizing single-nucleotide polymorphism (SNP) markers as in (93). The SNP markers located at the boundaries of the chromosome domains were chosen based on data from WormBase (WS231) and (54).The SNP markers and primers used are listed in (71). PCR and restriction digests of single embryo lysates were performed as described in (94, 95). Statistical analysis was performed using the two-tailed Fisher’s Exact test and Chi square test, 95% C.I., as in (96, 97).

## Acknowledgements

We thank Anne Villenueve and Abby Dernburg for generously providing antibodies and the Caenorhabditis Genetic Center for providing strains. We are grateful to Foxy Robinson, Merri-Grace Allard, Jaren Spotten, and Lorena Cruz-Gutierrez for help with embryonic lethality assays and construction of strains, and the Engebrecht and Colaiacovo labs for thoughtful discussions. We also thank MCB Light Imaging Facility Director, Michael Paddy for his help and patience with collection and processing of images and N. Silva and V. Jantsch for sharing unpublished data. This work was supported by National Institutes of Health (NIH) grant GM103860, UC Cancer Research Coordinating Committee CRR-17-426816 and Agricultural Experimental Station California-Davis grant *MCB-7237-H to J. Engebrecht, NIH grant T32GM0070377 to A.J. Deshong and K.S. Lawrence, NIH T32 CA10849 to P. M. Checchi, and NIH grants R01GM105853 and R01GM072551 to M. P. Colaiacovo.

